# Fetal stage melanopsin (OPN4) and GNAQ (Gαq) signaling regulates vascular development of the eye

**DOI:** 10.1101/537225

**Authors:** Shruti Vemaraju, Gowri Nayak, William E. Miller, David R. Copenhagen, Richard A. Lang

## Abstract

Maturation of sensory systems in mammals is regulated by appropriate sensory stimulation. Developmental refinement of the eye and visual system is regulated by light and visual stimulation. One compelling example is that fetal mouse pups deprived of light exhibit altered vascular development in their eyes. Previous work demonstrated that light activation of the photopigment melanopsin (*Opn4*), an atypical opsin expressed in intrinsically photosensitive retinal ganglion cells (ipRGCs), is crucial to normal vascular development. This suggested the unusual hypothesis that vascular development of the eye was regulated by ipRGC responses in the fetal eye by light that traveled through the body wall of the mother. Here, we test the requirement of OPN4 during fetal stages using genetic approaches. The G-protein GNAQ (Gαq) is a candidate mediator of melanopsin signaling. We show that ipRGC-specific deletion of *Gnaq* phenocopies both hyaloid and retinal vascular development of the *Opn4* null mouse. Furthermore, GNAQ gain-of-function in *Opn4*-expressing cells only during late gestation was sufficient to reverse the consequences for vascular development of either dark rearing or *Opn4* loss-of-function. We conclude that melanopsin-dependent signaling in the fetal mouse eye is necessary and sufficient for vascular maturation.

## Introduction

Melanopsin is a G-coupled receptor that, like other opsins^1–4^, initiates a signaling cascade when it captures a photon of light. Melanopsin was discovered in the melanophores of the frog *Xenopus laevis*^5^ but is now known to be present in most vertebrates^6^. In adult mice, melanopsin mediates circadian entrainment^7–10^ and the pupillary light reflex^9,11^. More recently, melanopsin was shown to be essential for negative phototaxis in neonates^12^ and for normal vascular development of the eye^13^.

In mammals, melanopsin is expressed in so-called intrinsically photosensitive retinal ganglion cells (ipRGCs). The axons of ipRGCs project to several regions of the brain, such as the suprachiasmatic nucleus (SCN)^14–18^ and the olivary pretectal nucleus^11,17,19–22^, that are consistent with the light-sensing physiological functions of melanopsin. It has been shown that melanopsin expressing cells are photo-sensitive as early as the day of birth (P1) in the mouse^23^. This is 9 days prior to the emergence of visual signaling from conventional photoreceptors, consistent with the acute action of melanopsin in the negative phototaxis reflex of mice at P5^12^. Several studies have demonstrated that ipRGCs express Gnaq/11 subfamily members^24–26^ and that melanopsin phototransduction is predominantly mediated by Gnaq/11 coupling^24,27,28^. Other reports suggest that melanopsin functions independent of Gnaq/11 both *in vivo*^26^ and *in vitro*^29–31^. The *in vivo* studies were performed in neonatal pups or adult animals, and the nature of melanopsin phototransduction in fetal retina remains to be explored.

Melanopsin is required for normal vascular development of the mouse eye^13^. Mice that are dark reared from E15 or that are *Opn4* loss-of-function show promiscuous retinal angiogenesis and hyaloid vessel persistence^13^. A model explaining melanopsin-dependent vascular development suggests that the first step is the suppression of retinal neuron numbers^13^. It is proposed that in *Opn4* null and dark reared mice, abnormally high numbers of retinal neurons induce a hypoxia response that results in high expression of vascular endothelial growth factor A, (VEGFA)^13^, a key regulator of developmental angiogenesis. In turn, high levels of VEGFA are proposed to cause promiscuous angiogenesis in the retina as well as to suppress regression of the hyaloid vessels^13^. One unusual outcome of this analysis was the proposal, based on staged dark rearing experiments, that the light response window for melanopsin-dependent vascular development occurs during gestation between E16 and E18^13^. This further implied that melanopsin in the fetal retina would be stimulated by light that traveled through the body wall of the dam^13^. Measured levels of light within the visceral cavity of adult female mice were consistent with this possibility given the threshold photo-sensitivity of ipRGCs^23,32,33^.

In the current study we have used experimental strategies to address the role of melanopsin signaling during fetal mouse stages in regulating vascular development in the eye. We show that gain- and loss-of-function modulation of GNAQ (Gαq) in *Opn4*-expressing cells results in changes in vascular development. Importantly, *Gnaq* loss-of-function phenocopies loss-of-function for *Opn4* while GNAQ gain-of-function restricted to fetal *Opn4*-expressing cells reverses the vascular consequences of dark rearing and *Opn4* loss-of-function. These data make a strong case that melanopsin function via GNAQ signaling during fetal stages is necessary and sufficient for vascular maturation in the eye. Given the OPN4 gene is expressed at gestational week 9 in human eyes^34^, these findings also raise the very interesting possibility that this same pathway may be conserved in the human fetus.

## Results

### *Gnaq* loss-of-function in *Opn4*-expressing cells phenocopies *Opn4* mutation

In order to employ manipulations of G-protein signaling to investigate fetal ipRGC vascular development responses, we first needed to establish which G-protein was involved. If ipRGCs employ GNAQ for melanopsin phototransduction, then a deletion of *Gnaq* in *Opn4*-expressing cells in mice should reproduce the vascular development phenotype of the *Opn4* null^13^. To test this, we combined the *Opn4^cre^* allele with *Gnaq^fl^* (officially *Gnaq^tm2Soff^*) but also included the germ-line loss-of-function *Gna11* allele because GNAQ and GNA11 frequently function together. To assess the consequences of deletion of GNAQ and GNA11 in *Opn4*-expressing cells, we generated *Opn4^cre^; Ai14, Gna11^−/−^; Gnaq^fl/fl^* mice and assessed deletion using immunofluorescence detection of GNAQ and GNA11. ipRGCs in the retina were identified by the expression of tdTomato from the *Ai14* cre reporter (Fig. 1A-E). Use of a single antibody that detects both GNAQ and GNA11 on cryosections from retina revealed that control ipRGCs showed membrane immunoreactivity (Fig. 1A-E), but almost all those from mice with combined *Gna11* and *Gnaq* mutations did not (Fig. 1F-H). A few ipRGCs retained GNAQ/11 labeling (Fig. 1G, H, red arrow) and probably represent cells that did not recombine both *Gnaq^fl^* alleles in response to *Opn4^cre^*. With this evidence for GNAQ/11 loss in ipRGCs, we analyzed the eyes of mutant mice for vascular anomalies.

**Figure 1.**
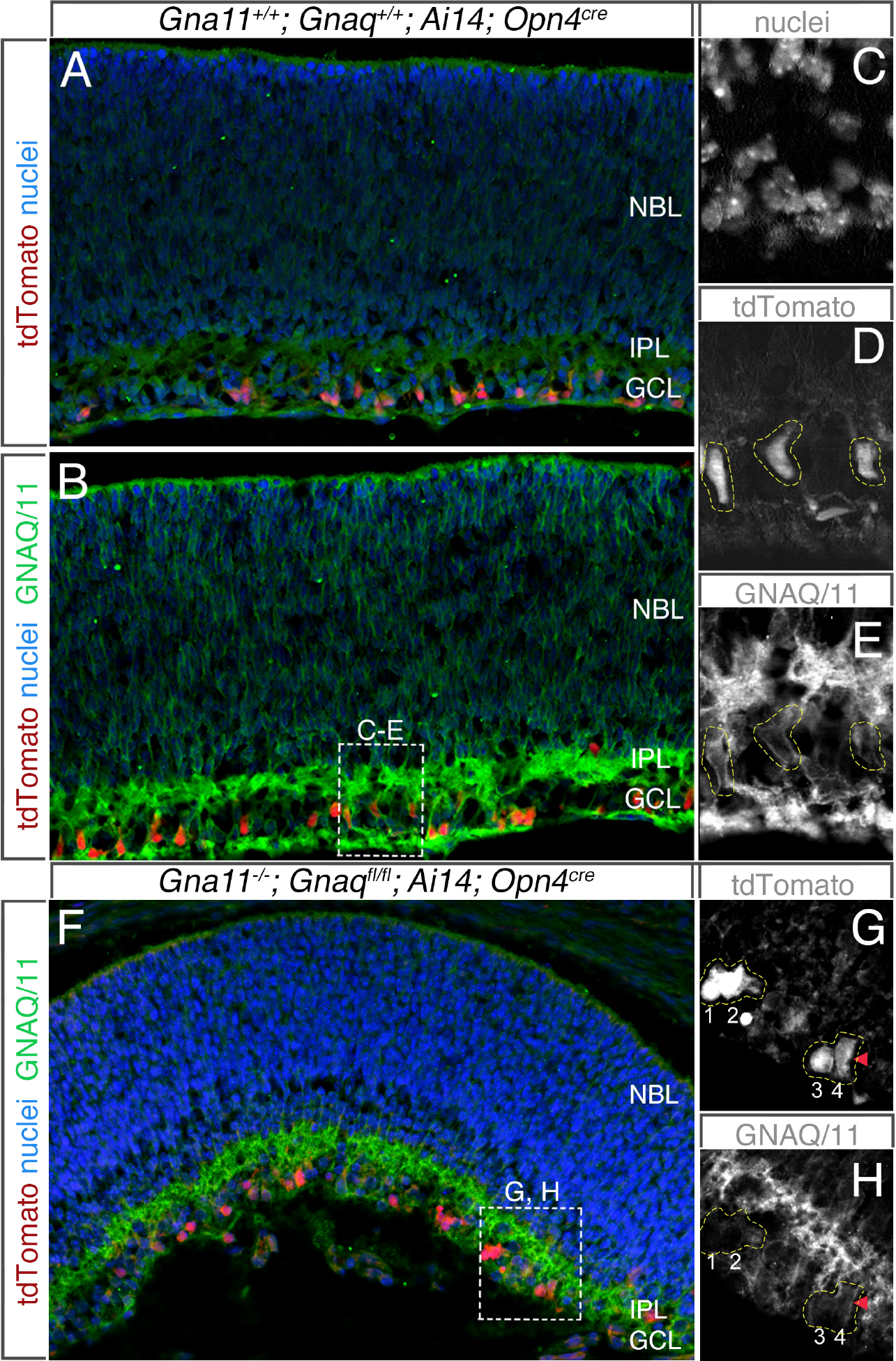
GNAQ loss-of-function in ipRGCs can be generated with *Opn4^cre^*. (A-E) Cryosections from P2, *Gna11^+/−^; Gnaq^+/+^; Ai14; Opn4^cre^* mice were labeled for nuclei (A, B, blue, C, grayscale), tdTomato (A, B, red, D, grayscale) and GNAQ/11 (B, green, E, grayscale, A no primary antibody control). C-E represent different channels from region demarcated in (B). In (E, yellow dashed lines), GNAQ/11 labeling is associated with the membrane of three ipRGCs according to the presence of tdTomato as shown in (D, yellow dashed lines). (F-H) Cryosections from E18, *Gna11^+/−^; Gnaq^fl/fl^; Ai14; Opn4^cre^* mice were labeled for nuclei (F, blue), tdTomato (F, red, G, grayscale) or for GNAQ (F, green, H, grayscale). G-H represent different channels from region demarcated in (F). In (H), the cells labeled 1-4 are ipRGCs according to the expression of tdTomato (G). Unlike ipRGCs in the control sections (D, E), cells 1 and 2 (G, H, yellow dashed line) do not show GNAQ/11 membrane labeling. Cells 3 and 4 (G, H, red arrowhead) show some GNAQ labeling and may therefore have escaped cre-mediated deletion of both *Gnaq^fl^* alleles. NBL neuroblast layer, IPL inner plexiform layer, GC ganglion layer

The hyaloid vessels are fetal blood vessels that reside transiently between lens and retina. These vessels undergo a maturational regression, presumably as an adaptation to clear the visual axis for high-acuity vision. In the mouse this occurs between postnatal (P) days 1 and 10^35^. Counts of vessel numbers at P8 provides a simple and reliable read-out for regression activity^13,36^. In P8 *Gnaq^fl/+^*; *Gna11^+/−^* control mice, we observed hyaloid vessel numbers consistent with a normal rate of regression (Fig. 2A-C)^13,36^. By contrast, *Opn4^cre^; Gnaq^fl/+^* and *Opn4^cre^; Gnaq^fl/fl^* both showed hyaloid persistence. Deleting both alleles of *Gnaq* produced the highest level of persistence suggesting a gene dose-dependence (Fig. 2A-C). The addition of *Gna11* germ-line homozygosity to *Gnaq* conditional deletion in ipRGCs did not further elevate hyaloid vessel number at P8 (Fig. 2C). These data suggest GNAQ is required in *Opn4*-expressing cells for activity of the pathway that controls hyaloid regression. The data also suggest that GNA11 is unlikely to be a significant contributor.

**Figure 2.**
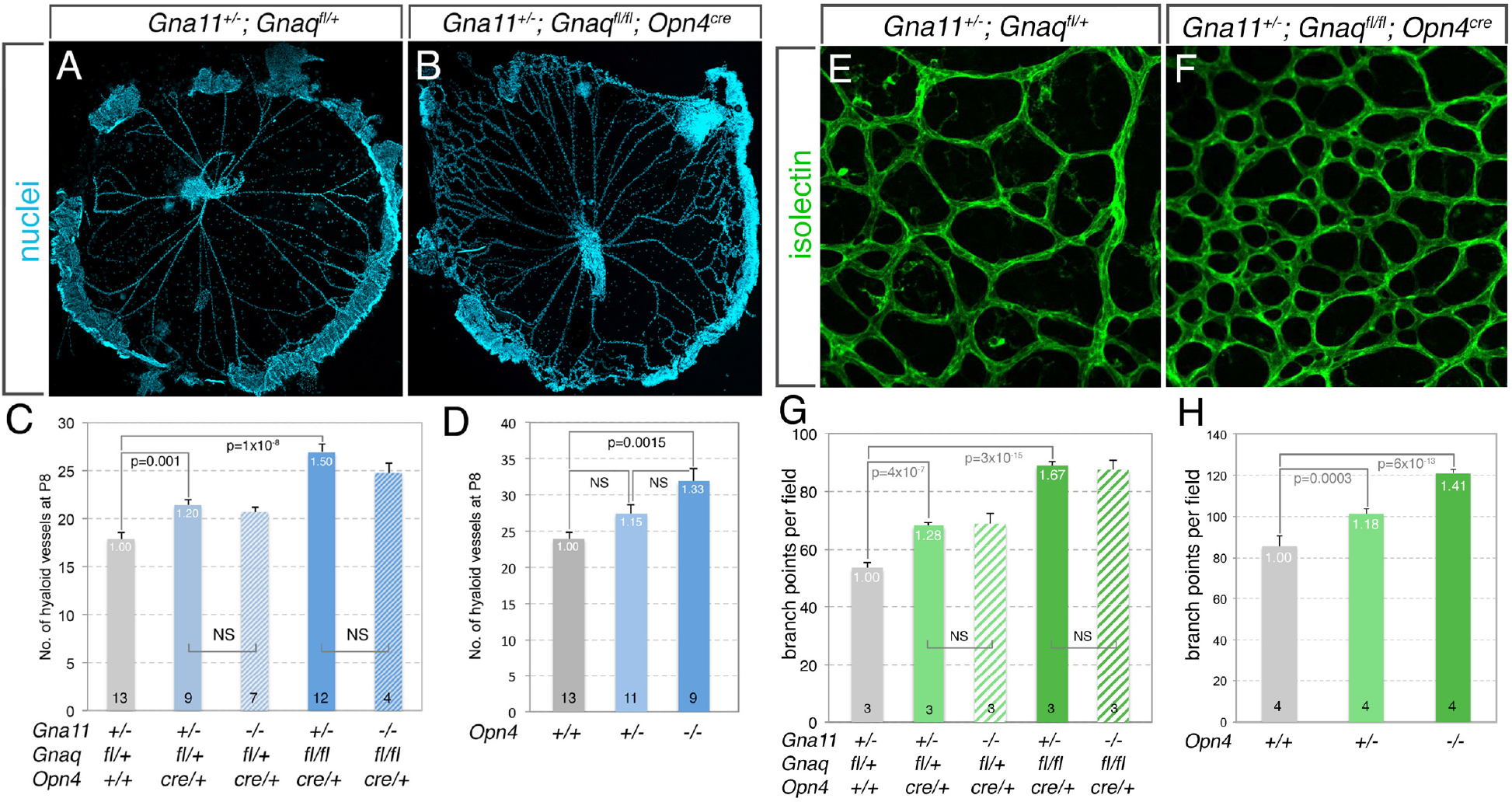
*Gnaq* loss-of-function in ipRGCs results in hyaloid vessel persistence and retinal vascular overgrowth. (A, B) Hoechst 33258-labeled hyaloid vessel preparations for control *Gna11^+/−^*; *Gnaq^fl/+^* (A) and experimental *Gna11^+/−^; Gnaq^fl/fl^; Opn4^cre^* (B) mice at P8. (C) Number of hyaloid vessels in P8 mice of the indicated genotypes. (D) For comparison with (C), number of hyaloid vessels in P8 *Opn4* wild type, heterozygote and homozygote mice. (E, F) Flat mount retina preparations labeled for vasculature with isolectin from control *Gna11^+/−^; Gnaq^+/fl^* (E) and experimental *Gna11^+/−^; Gnaq^fl/fl^; Opn4^cre^* (F) mice at P8. (G) Number of retinal vascular branch-points in P8 mice of the indicated genotypes. (H) Number of retinal vascular branch-points in P8 mice in *Opn4* wild type, heterozygote and homozygote mice. Data represented as mean ± S.E.M. Sample size is shown at the base of the each histogram and represents number of animals from multiple litters. p values calculated by Student’s T-test. p values as labeled, NS, not significant, for (C, D, G, H) white number at top of bars indicates the proportion of the control value for each panel.

As previously shown *Opn4* deletion results in persistent hyaloid vessels at P8^13^. For comparison purposes we also show here that *Opn4* homozygote null mice show a significant elevation in the number of hyaloid vessels (Fig. 2D). The absolute numbers of hyaloid vessels in control animals shown in Fig. 2C and 2D is distinct and this very likely reflects differences in strain background. Namely, the data in Fig. 2C are the result of a mixed background cross and that in Fig. 2D is C57BL/6 background. Any changes in hyaloid vessel number are thus best compared as a proportion of the value from littermate controls. When we express hyaloid vessel number for both the germline *Opn4* and conditional *Gnaq* mutants as a proportion of control (white numbers, Figs. 2C, D), we find that GNAQ loss-of-function produces a phenotype that is at least as severe as that for OPN4. This argues that for this vascular development pathway GNAQ activity can fully account for the signaling activity of OPN4.

During the postnatal period in mice, the retina becomes vascularized through a process of angiogenesis in part driven by VEGFA^37^. In the *Opn4* null mouse, retinal angiogenesis is promiscuous and the vascular networks show elevated density and ectopic sprouts^13^. For this reason we also assessed development of the retinal vasculature in the *Gna11^−^*, *Gnaq^fl^*, *Opn4^cre^* allelic series. Counts of branch-points revealed a gene dosage-dependent elevation in vascular density (Fig. 2E-G) but, as with the hyaloid vessel phenotype, no apparent contribution from GNA11. When we compared the phenotype severity for GNAQ and OPN4 loss-of-function (as proportion of control, white number Figs. 2G, H), as with the hyaloid vessel phenotype, we found that GNAQ can fully account for the activity of OPN4. Combined, the hyaloid vessel and retinal angiogenesis phenotypes of mice with GNAQ loss-of-function in *Opn4*-expressing cells suggest that OPN4 signaling for this vascular development pathway is coupled through GNAQ.

### Activation of GNAQ at fetal stages reverses the vascular consequences of *Opn4* loss-of-function

To further assess the relationship between light responses, OPN4, GNAQ and vascular development, we determined whether we could reverse the vascular consequences of dark-rearing or *Opn4* mutation by activating GNAQ signaling only in *Opn4*-expressing cells. We designed these experiments so that we could activate GNAQ only during late gestation and in fetal mice, not in the pregnant dam. This addressed the question of whether OPN4-dependent signaling responses occurred in fetal mice.

For these experiments, we employed a *ROSA26*-based allele expressing hM3Dq^38^, a DREADD (Designer Receptors Exclusively Activated by Designer Drugs) based on a G-coupled receptor that activates GNAQ in response to the small molecule ligand CNO^39^. By activating *R26-hM3Dq* with *Opn4^cre^* (Fig. 3A), we could obtain expression in *Opn4*-expressing cells and by injecting CNO into the dam in late gestation (Fig. 3B), we could restrict GNAQ activation to fetal stages. We also performed crosses that excluded the possibility of GNAQ activation in the dam. We used this experimental strategy to generate a series of control and experimental mice that were assessed for both hyaloid vessel regression and retinal vascular density.

**Figure 3.**
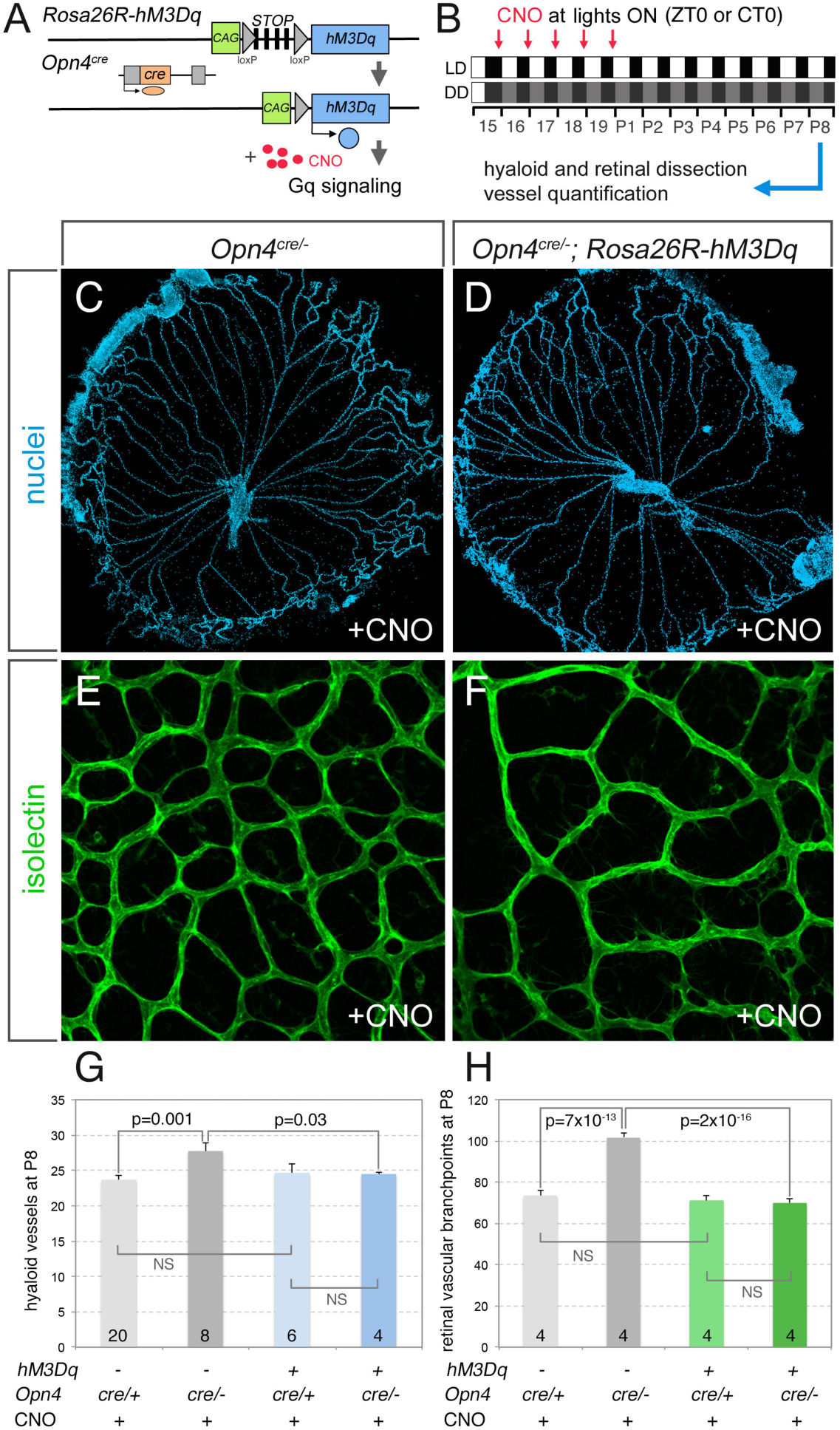
GNAQ gain-of-function in ipRGCs during gestation reverses the postnatal vascular consequences of *Opn4* mutation. (A) Schematic describing how the *R26-hM3Dq* and *Opn4^cre^* alleles can be combined with the artificial hM3Dq ligand CNO to give GNAQ signaling only in ipRGCs. (B) Experimental design for chemogenetic approach using hM3Dq/CNO. CNO ligand is injected into dam from E16 to P1 (red arrow) either in LD lighting cycle or after the transition to DD at E16 at lights ON (ZT0 for LD or subjective lights ON, CT0 for DD). At P8, hyaloid and retinal vessels are quantified. Hyaloid (C, D) and retinal (E, F) vessel preparations from *Opn4^cre/-^* (C,E) and *Opn4^cre/−^; hM3Dq* (D, F) mice that have received CNO from E16 to E19. (G) Quantification of P8 hyaloid vessel numbers in mice of the labeled genotypes and CNO exposure. (H) Quantification of P8 retinal vascular branch-points in mice of the labeled genotypes and CNO exposure. LD, normal lighting conditions, DD, constant darkness after E15. *cre*, cre recombinase coding region, *CAG*, chicken β actin promoter. p values as labeled; NS, not significant. Data represented as mean ± SEM. Sample size is shown at the base of each histogram and represents number of animals from three independent experiments. p values calculated using Student’s T-test.

For experiments to determine whether GNAQ activation could reverse the vascular consequences of *Opn4* loss-of-function, we crossed an *Opn4^cre/cre^*; *R26-hM3Dq* male with an *Opn4^+/−^* female. This provided appropriate control and experimental genotypes in fetal mice but excluded the possibility that the dam could respond to CNO. We then injected CNO daily from E16 to day of birth (P1) and harvested retina and hyaloid at P8 for quantification of vascular structures. As expected, *Opn4^cre/−^* pups showed hyaloid persistence at P8 (Fig. 3C, and the dark gray bar in G). By contrast, *Opn4^cre/−^*; *R26-hM3Dq* pups injected with CNO showed a normal number of hyaloid vessels (Fig. 3D, and the dark blue bar in G). Similarly, *Opn4^cre/−^* animals showed an abnormally elevated retinal vascular density at P8 (Fig. 3F, H) and this was reversed when *Opn4^cre/−^* was combined with *R26-hM3Dq* and gestational CNO (Fig. 3F, H). These data show that for the vascular development pathways in the eye, GNAQ activity can substitute for OPN4 within ipRGCs. The experiment also makes a strong case that this vascular development pathway is dependent on the activity of melanopsin at fetal stages.

### Activation of GNAQ in ipRGCs at fetal stages reverses the vascular consequences of dark rearing

As a second step in this analysis, we determined whether fetal stage GNAQ activation could reverse the vascular consequences of dark-rearing (DD, Fig. 3B). For this analysis, we crossed a male *Opn4^cre/cre^* homozygote with a female *R26-hM3Dq* heterozygote. This meant that half the fetal mice had expression activation of hM3Dq in melanopsin-expressing neurons, but importantly, the dam did not carry *Opn4^cre^* and therefore could not activate expression of *R26-hM3Dq.* Thus, any response to the CNO ligand could only be attributed to fetal *Opn4*-expressing cells. When we performed the analysis, we found that CNO injected into pregnant female mice from E15 through E19 reversed the vascular consequences of dark rearing (Fig. 4A-H). As expected, DD conditions resulted in significantly elevated hyaloid vessel numbers at P8 (Fig. 4A, C, D). In the absence of late gestational injections of CNO the elevated number of hyaloid vessels was not significantly changed with or without hM3Dq (Fig. 4C, D). However, when CNO was injected from E15 to E19 into DD, *Opn4^cre^*; *R26-hM3Dq* mice, the normally elevated number of hyaloid vessels remained at the level observed in the LD control (compare Fig. 4E, blue bar with 4C, both bars). When we quantified retinal vascular branch-points (Fig. 4F, G, H), we observed a very similar pattern where *Opn4^cre^*, *R26-hM3Dq*, and gestational CNO were all required to reverse the retinal vascular overgrowth of DD to control levels (Fig. 4H). These data indicate that activation of GNAQ signaling in *Opn4*-expressing cells can substitute for light in late gestation and give normal regulation of vascular development.

**Figure 4.**
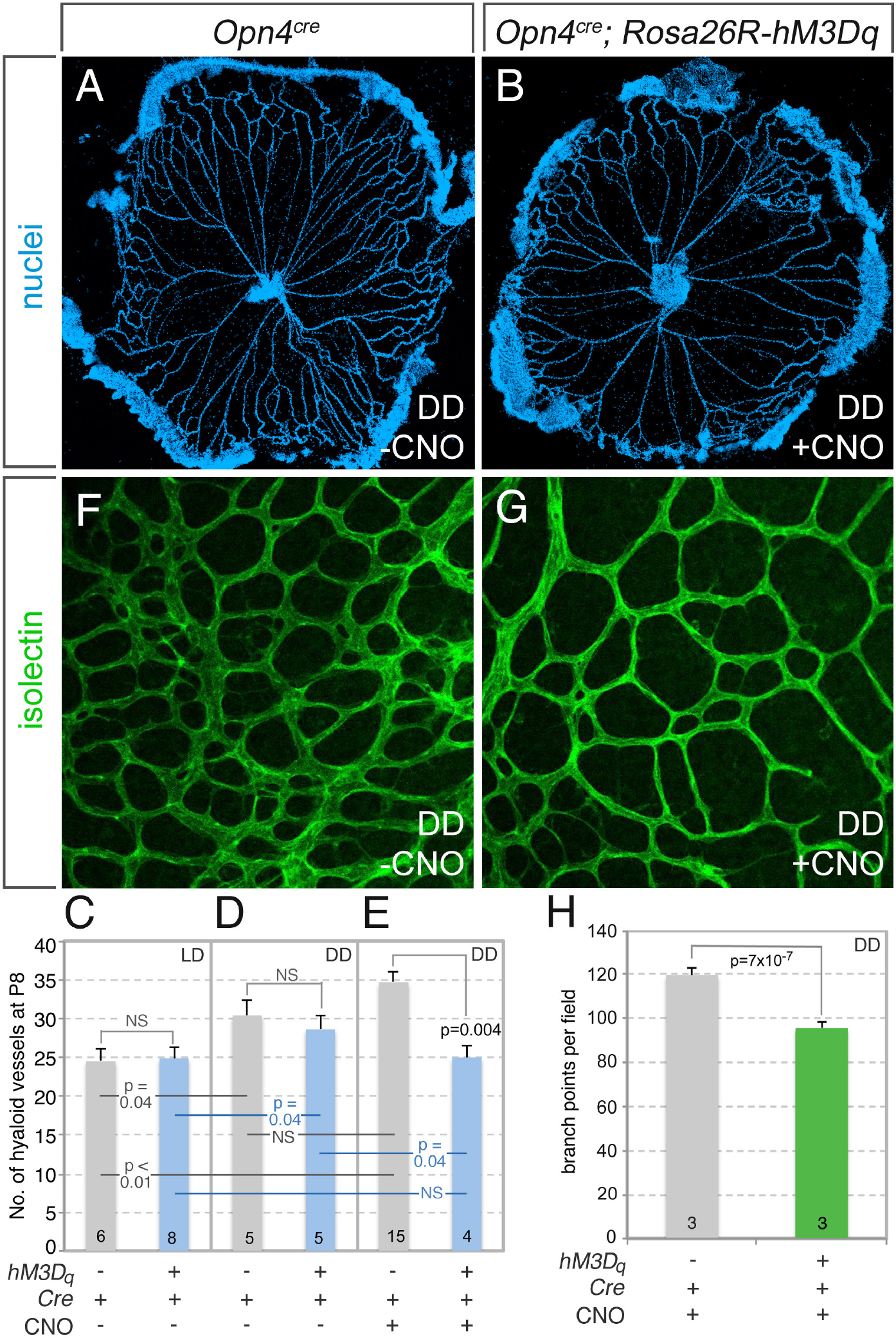
GNAQ gain-of-function in ipRGCs during gestation reverses the postnatal vascular consequences of dark-rearing. (A, B) Hyaloid vessel preparations at P8 for *Opn4^cre^* (A) and *Opn4^cre^; hM3Dq* (B) mice where dam was moved to DD at E15 and injected with CNO from E16 to day of birth P1 (see Fig. 3b for experimental design). (C-E) Quantification of P8 hyaloid vessel numbers in mice of the labeled genotypes and CNO exposure. (F, G) Retinal vessel preparations at P8 for *Opn4^cre^* (F) and *Opn4^cre^; hM3Dq* (G) mice moved to DD as above. (H) Quantification of P8 retinal vascular branch-points in mice of the labeled genotypes and CNO exposure. LD, normal lighting conditions, DD, constant darkness after E15. p values as labeled; NS, not significant. Data represented as mean ± S.E.M. Sample size is shown at the base of each histogram bar and represents number of animals from three independent experiments. p values calculated using Student’s T-test.

## Discussion

Here we have combined experimental strategies to address the question of whether melanopsin activation during fetal stages can regulate vascular development in the mouse eye. Within the retina, OPN4 expression is restricted to a subset of retinal ganglion cells (RGCs) that are, in adult mice, intrinsically photosensitive^9,14,40^. *Opn4* is expressed as early as E10.5^34^ and OPN4 protein is detected in the retina around E15^41^ in mouse embryos. Like other opsins, melanopsin (aka Opsin 4, OPN4) is a G-coupled receptor^29,42^ and functions predominantly via Gnaq/11 coupling^24,27,28^. Few studies have suggested alternate G-protein coupling for melanopsin in both *in vivo*^26^ and in expression systems^29–31^. Current data on melanopsin G-protein coupling is from adult or neonatal mice. We focused our efforts on late gestational ages and our data indicates that melanopsin in the fetal retina uses conventional Gnaq/11-coupling for ocular vascular development.

In the first step of this analysis, we determined whether the loss of GNAQ function in *Opn4*-expressing cells produced the same phenotype as a melanopsin (OPN4) loss-of-function. We show mice with the *Opn4^cre^; Gnaq^fl/fl^* genotype had the retinal vascular overgrowth and hyaloid vessel persistence characteristic of the *Opn4* null and of dark rearing from late gestation^13^. Furthermore, the addition of germ-line *Gna11* homozygosity to *Gnaq* conditional deletion from *Opn4*-expressing cells did not exacerbate vascular phenotypes. These data argue that GNAQ, and not GNA11, mediates melanopsin-dependent vascular development. As a final step in this series of experiments we determined whether GNAQ gain of function was sufficient to reverse the vascular phenotypes that result from dark rearing or from *Opn4* mutation. We tested whether GNAQ activation in *Opn4*-expressing cells only during the fetal stages of mouse development was sufficient to reverse the vascular changes that normally result from dark rearing or *Opn4* loss-of-function. Our findings showed that when GNAQ gain-of-function was restricted to *Opn4*-expressing cells in late mouse gestation, this was sufficient to overcome the absence of light or the absence of OPN4 for the vascular development pathway.

These data thus make a strong case that if the vascular development of a mouse eye is to be normal, GNAQ must be activated in *Opn4*-expressing cells from E16 to E19 of gestation. The ability to revert the *Opn4* loss-of-function phenotype with conditional activation of GNAQ also suggests that the activity of OPN4 and GNAQ in *Opn4*-expressing cells are directly related. The ability to revert the vascular consequences of dark rearing also indicate that the fetal retina normally responds to environmental light. We show that the fetal light response pathway regulating vascular development requires GNAQ but not GNA11 (Fig. 2). Involvement of GNA14 in the fetal ipRGC response remains a possibility though this is unlikely given the quantitative correspondence between *Opn4* and *Gnaq* loss-of-function.

Multiple members of the *Gnaq/11* subfamily, including *Gna14, Gnaq* and *Gna11*, can participate in melanopsin phototransduction *in vivo*^24,27,28^. However, a recent study showed that *Opn4^cre^; Gna11^−/−^; Gnaq^fl/fl^* mice like those used here, many of the photic responses ascribed to melanopsin remain normal^26^. The present analysis, showing that GNA11 does not contribute to vascular development responses, is consistent with the documented lack of a multielectrode array response in the neonatal retina of *Gna11; Gna14* null mice^26^. However, the *Gnaq* loss-of-function phenotype, coupled to the rescue of the OPN4 loss-of-function phenotype by GNAQ gain-of-function, makes a compelling case that GNAQ is an important signaling mediator of OPN4 activity in fetal stages and is consistent with the observation of GNAQ presence in all M1-5 ipRGCs in adult mice^24^. One possible explanation for these apparently discrepant findings is that different functions of melanopsin are mediated by distinct types of signaling response and are likely to be age-specific. One scenario is that perhaps the vascular development pathway assessed here relies on the activity of GNAQ while the pupillary reflex and circadian photoentrainment rely on other G-proteins that are expressed at a later stage of ipRGC development.

The data presented here make a strong case that the eye of a fetal mouse uses light as normal developmental cue. Though we currently do not understand the biological rationale with precision, it is quite likely that the fetal eye light response prepares the eye for visual function. Certainly, vascular development of the eye must be closely regulated for normal vision but it is also likely that other responses to fetal OPN4 stimulation will be identified. OPN4 is highly conserved within the vertebrates both in primary sequence and function. This raises the fascinating possibility that the human fetal eye is also light responsive in a way that regulates vascular development. This idea has received support from a recent clinical study that associates first trimester fetal light exposure with severe forms of retinopathy of prematurity, an ocular blood vessel overgrowth of the premature infant^43^.

## Acknowledgements

We are indebted to Dr. Ute Hochgeschwender for providing *R26-LSL-Gq-DREADD* mice prior to publication. We are also grateful to Dr. Ute Hochgeschwender and Dr. Shawnta Y. Chaney for critiquing this MS. We thank Mr. Paul Speeg for excellent technical assistance. We acknowledge funding from the National Eye Institute (EY021636-01 to RAL and DRC) in support of this work.

## Materials and Methods

### Animals

All animal experimentation was carried out using protocols approved by the Institutional Animal Care and Use Committee at Cincinnati Children’s Hospital Medical Center. Mouse lines used in this study have been previously described: *Opn4^cre^*(ref^44^) *Opn4^−^* (ref^10^), *Ai14* (JAX #007914), *Gnaq^fl^* (ref^45^), *Gna11* (ref^45^) and *R26-LSL-Gq-DREADD* (ref^38^). Genotyping information is described in the cited publications. Mice were housed in a 12:12 hour light-dark (LD) cycle. For dark-rearing experiments, pregnant dam was moved to constant darkness (DD) after lights OFF at E15. Clozapine-N-oxide (CNO, Sigma #CO832-5MG) stock solution was prepared at 10 mg/ml in 100% DMSO and injected into the dam at 1 mg/kg body weight to activate *Gq-DREADD* (*hM3Dq*). Dam was injected daily at lights ON in LD cycle or at subjective lights ON for animals in DD from embryonic day E16 to day of birth. For DD animals, injections were performed under dim red light. Littermate controls were used for all experiments to accommodate for differences in mouse strains.

### Hyaloid and retinal labeling

Hyaloid vessel and retinal flat-mount preparations have been previously described (Rao 2013). Hyaloid preparations and retinae were labeled with Hoechst 33258 and isolectin (1:500, Thermo Fisher Scientific #I21411), respectively. Hyaloid vessel quantification has been described previously (Rao 2013). Retinal vessel density was quantified by counting vessel junctions using ImageJ for at least ten 320μm^2^ fields per retina. We used paired T-test to assess statistical significance. In the G-protein immunohistochemical study, retinae for cryosections were fixed at room temperature for 2 hours in 4% PFA and then cryopreserved in 30% sucrose and sectioned at 10 μm followed by antigen retrieval with ice cold sodium citrate buffer for 10 minutes. Sections and flat-mounts were incubated in anti-GNAQ/GNA11 (1:100, Santa Cruz sc-392) at 4°C for 1-3 days followed by secondary antibody (1:500, Thermo Fisher Scientific #A-21206) overnight at 4°C.

Author contributions
DRC and RAL provided project leadership, designed experiments and supervised experimental work. RAL, DRC and SV wrote the manuscript. WEM provided crucial reagents and advice. SV and GN performed experimentation and analysis.

